# Modelling the impact of clot fragmentation on the microcirculation after thrombectomy

**DOI:** 10.1101/2020.11.30.403808

**Authors:** Wahbi K. El-Bouri, Andrew MacGowan, Tamás I. Józsa, Matthew J. Gounis, Stephen J. Payne

## Abstract

1

Many ischaemic stroke patients who have a mechanical removal of their clot (thrombectomy) do not get reperfusion of tissue despite the thrombus being removed. One hypothesis for this ‘no-reperfusion’ phenomenon is micro-emboli fragmenting off the large clot during thrombectomy and occluding smaller blood vessels downstream of the clot location. This is impossible to observe in-vivo and so we here develop an in-silico model based on in-vitro experiments to model the effect of micro-emboli on brain tissue. Through in-vitro experiments we obtain, under a variety of clot consistencies and thrombectomy techniques, micro-emboli distributions post-thrombectomy. Blood flow through the microcirculation is modelled for statistically accurate voxels of brain microvasculature including penetrating arterioles and capillary beds. A novel micro-emboli algorithm, informed by the experimental data, is used to simulate the impact of micro-emboli successively entering the penetrating arterioles and the capillary bed. Scaled-up blood flow parameters – permeability and coupling coefficients – are calculated under various conditions. We find that capillary beds are more susceptible to occlusions than the penetrating arterioles with a 4x greater drop in permeability per volume of vessel occluded. Individual microvascular geometries determine robustness to micro-emboli. Hard clot fragmentation leads to larger micro-emboli and larger drops in blood flow for a given number of micro-emboli. Thrombectomy technique has a large impact on clot fragmentation and hence occlusions in the microvasculature. As such, in-silico modelling of mechanical thrombectomy predicts that clot specific factors, interventional technique, and microvascular geometry strongly influence reperfusion of the brain. Micro-emboli are likely contributory to the phenomenon of no-reperfusion following successful removal of a major clot.

**Author summary:** After an ischaemic stroke - one where a clot blocks a major artery in the brain - patients can undergo a procedure where the clot is removed mechanically with a stent - a thrombectomy. This reopens the blocked vessel, yet some patients don’t achieve blood flow returning to their tissue downstream. One hypothesis for this phenomenon is that the clot fragments into smaller clots (called micro-emboli) which block smaller vessels downstream. However, this can’t be measured in patients due to the inability of clinical imaging resolving the micro-scale. We therefore develop a computational model here, based on experimental thrombectomy data, to quantify the impact of micro-emboli on blood flow in the brain after the removal of a clot. With this model, we found that micro-emboli are a likely contributor to the no-reflow phenomenon after a thrombectomy. Individual blood vessel geometries, clot composition, and thrombectomy technique all impacted the effect of micro-emboli on blood flow and should be taken into consideration to minimise the impact of micro-emboli in the brain. Furthermore, the computational model developed here allows us to now build large-scale models of blood flow in the brain, and hence simulate stroke and the impact of micro-emboli on the entire brain.

## 3 Introduction

Stroke is the second largest cause of mortality and of adult disability globally [1]. Ischaemic stroke – the physical occlusion of a cerebral artery by a blood clot (embolus) restricting blood supply to a part of the brain – is the most common form of stroke, occurring in approximately 85% of strokes [1]. Time is critical in stroke, with every hour of delay without treatment causing a loss of neurons equivalent to 3-4 years of ageing [2] and a 6% reduction in positive outcome once the patient is treated [3].

Since 2015, the use of intra-arterial thrombectomy – the mechanical removal of a clot via a stent – has become the standard of care for patients with acute ischaemic stroke caused by large vessel occlusion [4,5]. Despite this, many patients do not recover full perfusion in their tissue downstream of the recanalized vessel [6]. This observation, commonly known as the “no-reflow” or “no-reperfusion” phenomenon, has been documented in numerous studies involving both animals and humans [7–12]. Hypotheses for this phenomenon include: changes in the ultrastructure of the microvasculature and spontaneous blood clotting [7], capillary stalling and leukocyte adhesion [10,13–15], breakdown of the blood-brain barrier leading to tissue swelling and vessel collapse [12,16,17], vasoconstriction of the capillary vessels due to pericyte death [10,18,19], oxidative stress and inflammatory responses [11], and micro-emboli fragments post-recanalization blocking micro-vessels downstream of the clot [8,20].

One of the above hypotheses for the no-reflow phenomenon – a micro-emboli “shower” fragmenting off the clot – is of interest due to the clear clinical treatment pathway via thrombolysis. There is indirect evidence that micro-emboli can fragment off clots and lead to worse patient outcomes. Patients who undergo endarterectomy (surgical removal of a plaque in the vasculature) with stenting in the carotid artery are 3 times as likely to have lesions appear in diffusion weighted MRI scans, and have non-disabling strokes, than those who only had an endarterectomy [21]. There is therefore a potential link between clot fragmentation and the potential for microvascular occlusions and worse patient outcomes.

As part of the INSIST (*IN-silico clinical trials for treatment of acute Ischaemic STroke*) consortium (www.insist-h2020.eu) we are developing full brain models of blood flow and oxygen transport in health, after an ischaemic stroke, and post-thrombectomy/thrombolysis [22]. The goal of INSIST is to advance in-silico clinical stroke trials for biomedical products for the treatment of ischaemic stroke. As such, an accurate model of clot fragmentation during thrombectomy is required to simulate the effect of micro-emboli on reperfusion post-recanalization. This will help in deducing whether micro-emboli are a major reason for no-reperfusion.

Due to the intrinsically microvascular nature of the no-reflow phenomenon, it is impossible to assess the impact of micro-emboli showers in humans, with most microinfarcts being < 1 mm in size making in-vivo detection of them challenging [23]. However, silicone in-vitro models of human cerebral vasculatures have been used to quantify how a clot fragments during thrombectomy using a range of removal techniques and different clot consistencies [20,24]. Most other experimental works analysing the impact of occlusions on the microvasculature come from mouse models [25–28]. Recently, an in-silico study on a mouse cortex microvascular network found that the severity of micro-strokes in the capillary bed were heavily influenced by the local network topology, with decreases in flow of up to 80% in the local vicinity of the capillary [29].

On the macro-scale, full organ models of blood flow in the brain have been developed [30,31]. These models treat the microvasculature in the brain as a continuous porous medium, effectively smoothing out the local topology and variations in blood flow. Crucially, these models are parameterized by simulations of blood flow in the microvasculature [32]. As such, in order to achieve the aim of simulating a micro-emboli shower in the microvasculature and to determine its effect on the large-scale blood flow, a multi-scale model of micro-occlusions will be developed here. This model will link the blood flow in the microvasculature to the parameters used to model full brain blood flow and the effect of thrombectomy on the microcirculation, both at the penetrating vessel scale and the capillary scale. To the best of our knowledge no models of micro-emboli occlusions have been developed for post-thrombectomy patients.

This paper then seeks to answer, through modelling informed by in-vitro experiments, the following questions: To what extent is clot fragmentation responsible for the post-thrombectomy no-reflow phenomenon? Can the changes in the blood flow modelling parameters due to micro-emboli showers be accurately quantified such that they can be used in the full organ models? What impact do thrombectomy technique and clot consistency have on downstream blood flow post-thrombectomy?

## 4 Materials and methods

### 4.1 Modelling framework

In order to investigate the effects of micro-emboli on the cerebral microvasculature, a modelling framework must first be developed. There are two main ways in which the cerebral microvasculature can be modelled – either as a network of individual vessels [35–37], or as an averaged porous medium [30,31,33,38]. We model the microvasculature as a porous medium in order to allow large regions of the brain to be modelled computationally feasibly.

The microvasculature will be parameterised following on from previous research developed by our group [32,33,39]. Firstly, statistical models are developed which match morphological properties of the human cerebrovascular network [40–43]. The microvasculature is then split into 2 different spatial scales – the capillary bed, and the penetrating arterioles. In order to scale up the microvasculature to larger regions, the capillary bed blood flow is parameterised with a permeability, as are the penetrating arterioles. The flow from the penetrating arterioles to the capillary bed is parameterised through coupling coefficients – which represent the conductance of the pre-capillary arterioles. The parameterisation method is explained in Section 4.7. Example networks of the different microvascular scales and their coupling can be seen in Fig 1.

**Fig 1.**
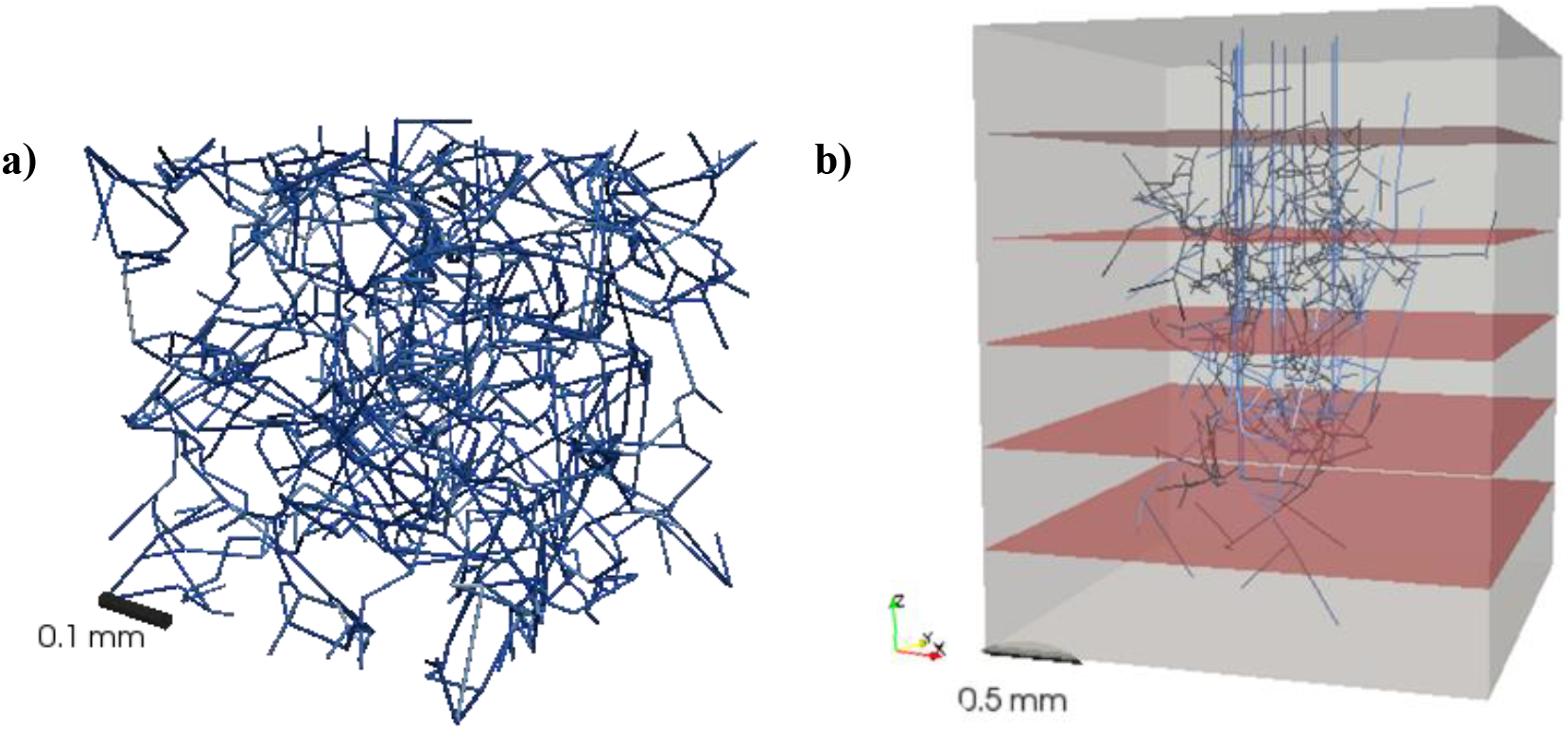
a) An example of a statistically accurate capillary network, b) An example of a voxel used to simulate flow through the penetrating arterioles. The grey region represents the homogenized capillary bed to which the terminal vessels of the trees couple. The red planes denote the dividing planes between the 6 layers of the voxel over which the blood flow characteristics will be calculated.

### 4.2 In-vitro emboli quantification

Previous in-vitro studies generated micro-emboli distributions from thrombectomy procedures. In brief, a silicone cerebrovascular model built from clinical imaging data was used in a flow loop with a peristaltic pump to simulate blood flow through the cerebral vasculature [24,44]. A thrombus was injected into the cerbrovascular replica to form a middle cerebral artery (MCA) occlusion. The thrombus was either defined as a soft or hard clot, and validated for bulk mechanical properties based on clinical specimens [45]. Four different clot removal techniques (thrombectomy) were then deployed to remove the clot from the MCA. Specifics on the 4 thrombectomy techniques – abbreviated as ADAPT, Solumbra, BGC, and GC at cervical ICA – can be found in Chueh et al. [20]. The number and size distribution of the clot fragments that break off the thrombus during thrombectomy are quantified using a Coulter counter (Beckman Multisizer 4, Brea, California). Each removal technique is repeated 8 times for each clot type, resulting in 64 sets of clot fragment data.

When sampling from this data for our in-silico models the 8 experimental micro-emboli density distributions (for a given removal technique and clot type) are averaged such that for any given technique and clot type there is only one average set of micro-emboli results relating number density and particle size. The smallest clot measurable, due to the aperture used in the Coulter counter, was 8.034 μm in diameter, with smaller clots binned under this size.

### 4.3 Simulating blood flow in the microvasculature

Prior work has shown how blood flow can be simulated in a coupled model of the penetrating vessels and a porous representation of the capillary bed [33]. This model is adapted here such that it can be used to determine the permeability of the penetrating arterioles in health and with micro-emboli occluding the vessels.

An example voxel, where the penetrating arterioles are coupled to the capillary bed, is shown in Fig 1b. Flow through the penetrating vessels is assumed to be steady-state, non-pulsatile (Womersley number << 1) and fully developed, hence approximated by Poiseuille flow:

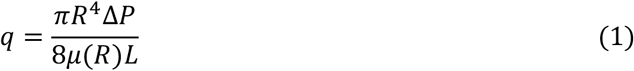

where *q* is the blood flow through the vessel, *R* is the radius of the vessel, Δ*P* is the pressure drop across that vessel, *L* is the length of the vessel, and *μ*(*R*) is the apparent viscosity of blood in the vessel correcting for the Fåhræus-Lindqvist effect with a constant discharge haematocrit of 0.45 [46].

As flow is conserved at each bifurcation node *i*, the net flow at each node in the tree is zero except for the one inlet node and the multiple terminal nodes that end in the capillary bed. Therefore, the flow through each internal node for *n* nodes can be written as:

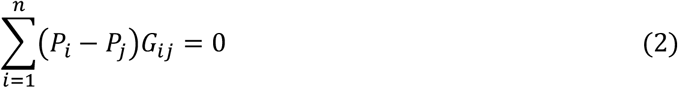

where *G*_*ij*_ is the flow conductance between nodes *i* and *j* and is defined as 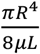 as in equation (1). The conductance is zero unless there is a vessel between nodes *i* and *j*. *P*_*i*_ and *P*_*j*_ are the pressures at nodes *i* and *j* respectively. The inlet flow and terminal flows are handled by the boundary conditions at the inlet to the arteriole and at the coupling points to the capillary bed. This set of *n* equations can be written in matrix form and solved as in Su et al. [47].

The capillary bed can be modelled as a porous medium using a volume-averaged form of Darcy’s law [32,48]:

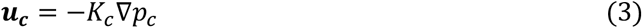

where *u*_*c*_ is the volume averaged capillary velocity, *K*_*c*_ is the permeability (as the permeability is isotropic this can be treated as a scalar), and ∇*p*_*c*_ is the pressure gradient across the capillary bed. The permeability *K*_*c*_ encapsulates the micro-scale geometry of the capillary bed in one averaged parameter, hence allowing for large-scale regions to be modelled computationally efficiently. This equation can be combined with mass conservation to give:

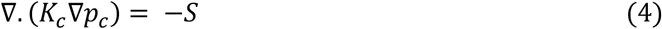

where *S* is the volumetric source/sink term [m^3^ s^−1^ m^−3^] that comes from the terminal nodes of the penetrating vessels. The healthy capillary permeability value has previously been found be 4.28×10^−4^ mm^3^ s^−1^ kg^−1^ [32]. Equation (4) is the Poisson equation that can be solved for capillary pressure over an appropriate 3D domain.

### 4.4 Coupling penetrating vessels and capillary bed

The coupling between the penetrating vessel terminal nodes and the porous capillary bed has a mathematical singularity at the point of coupling between 1D flow and 3D tissue. Previous models have approximated the Dirac delta singularity with a function that is spatially variant yet maintains mass conservation [30]; have approximated the pressure drop in the vicinity of the coupling point to deal with the strong pressure gradients that appear there [49]; or have simply taken the discretised element nearest the terminal node as the outlet region [33,50]. Here, we use a spatially variant function to approximate the volumetric source terms [30]. A shape function is introduced that replaces the Dirac delta singularity with an exponential in the vicinity of the source. The shape function, *η*, is defined as

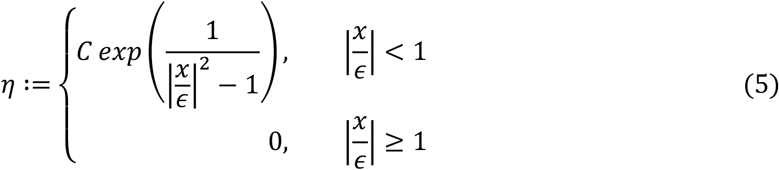

where *x* is the distance measured from the source/sink node and ε is the radius of the sphere over which the flow will be distributed from the arteriole source to the capillary bed (here set as 15 μm). *C* is a constant that acts to normalise *η* to ensure mass conservation across the capillary bed elements coupled to the terminal tree node. Therefore, the volumetric source term for each finite element within ∈ radius of the terminal node will be

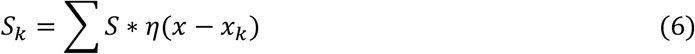

where *S*_*k*_ is the volumetric source term for one element within ε radius of the terminal node and *x*_*k*_ is the coordinate of the terminal node. A summation is used to account for the potential that multiple terminal nodes may feed an individual element. As the shape function *η* conserves mass, ∫ *η*(*x*)*dx* = 1 and hence ∑_*k*_ *S*_*k*_ = *S*.

### 4.5 Solving for blood flow under healthy conditions

The voxel size chosen to simulate the flow is 2 × 2 × 2.5 mm (on the order of an MRI voxel), with 8 arterioles /mm^2^ penetrating into the voxel [51]. This voxel was situated in a capillary bed of 2.25 × 2.25 × 2.5 mm to avoid terminal nodes being at the face of the voxel (Fig 1b). As the parameterisation of micro-emboli occluding the penetrating arterioles was primarily of interest here, the venules were not included in the simulation. As well as this, the permeability of the capillary bed was chosen such that the maximum possible flow could be obtained through the arterioles (in this case *K*_*c*_ was set to 4.2×10^−5^ mm^3^ s kg^−1^). The arterioles have a mean diameter of 20 μm ± 5 μm, with a mean length of 1.25 mm ± mm. An inlet pressure of 12.3 kPa is imposed at the tree inlets [52]. Dirichlet boundary conditions were imposed at the top and bottom of the voxel with *P*_*top*_ = 12.3 *kPa* at *z* = 0 and *P*_*bottom*_ = 8 *kPa* at *z* = 2.5 *mm*, with periodic boundary conditions on the 4 other faces.

The finite element method is used to simulate the capillary bed partial differential equation using FEniCS, an open source finite element solver [53]. Equation (4) is solved with the parameters and boundary conditions using 870,000 linear Lagrangian elements to obtain the pressure in the capillary bed. The equations above were solved iteratively. The pseudo-algorithm below details the solution steps:

#### Flow Solver Algorithm

1. Set Dirichlet boundary conditions on the top and bottom of the voxel and periodic boundary conditions on the 4 sides;
2. For all tree outlets coupled to capillary bed, determine elements in mesh coupled to the outlet node, and determine their *η* weightings using equation (5);
3. Set desired inlet pressures for penetrating vessels and set initial terminal outlet pressures = 0
4. Solve for the pressure in the trees using equations (1) and (2), hence solve for the outlet flows from the trees into the capillary bed weighting the volumetric source terms using equations (5) and (6);
5. Set the volumetric source terms in equation (4) and solve for the capillary bed pressure using the finite element method;
6. Update penetrating vessel terminal node pressure values using an *η* weighted average of the capillary bed pressures related to the given terminal node;
7. Go to step 4 and repeat, unless a steady state is reached where pressures in terminal nodes and capillary bed have converged to a tolerance of 1×10^−3^ Pa;

### 4.6 Simulating micro-emboli post-thrombectomy

In order to simulate the effect of micro-emboli on the microvasculature, an algorithm is presented here that samples from the in-vitro experimental thrombectomy data and occludes vessels sequentially. The algorithm flowchart is schematically represented in Fig 2.

**Fig 2.**
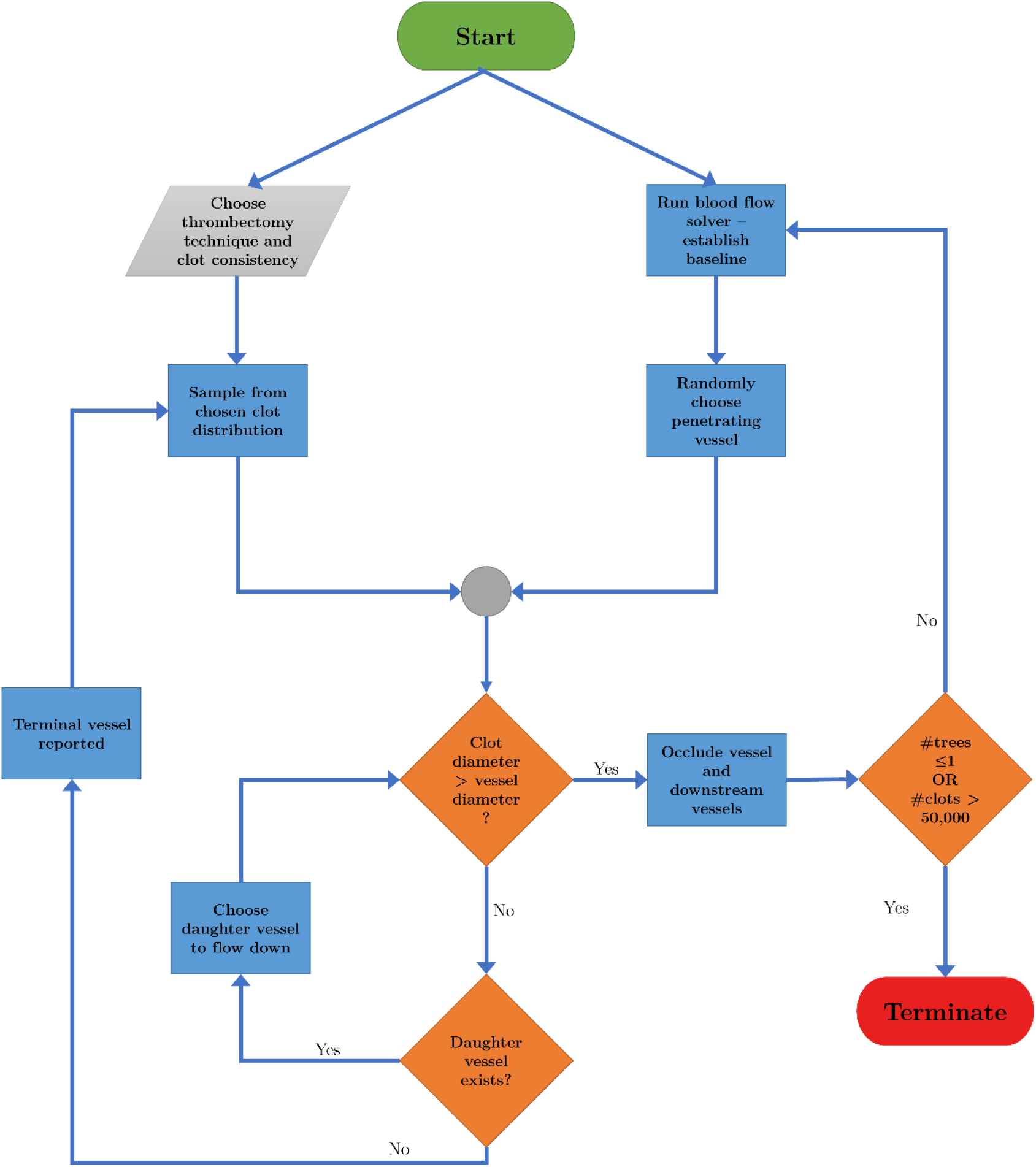
Flowchart representation of the micro-emboli occlusion algorithm

A clot technique and clot consistency are chosen from one of our 8 clot distributions as described in Section 3.2. The algorithm initially runs the healthy flow solver above to establish baseline healthy blood flow. A clot size is then sampled pseudo-randomly using Python’s inbuilt ‘random’ module from the experimental distributions derived from the in-vitro models [20].

A penetrating arteriole is randomly chosen on the pial surface through which the clot is sent through. The clot is assumed to be perfectly spherical. A check is then made to determine whether the clot is larger than the diameter of the vessel it is entering. If the clot is smaller than the vessel, the clot is assumed to travel until the next bifurcation. At the bifurcation, the clot enters one of the two daughter branches based on a probability that is linearly weighted by the blood flow in the daughter branches. For example, if one daughter vessel has 3 times as much flow as the other daughter vessel, the clot has a 75% chance of going down the vessel with more flow.

The clot continues to travel through vessels if its diameter is smaller than the diameter of the vessel it is entering. If the clot enters a vessel with a smaller diameter, it is assumed that the clot completely occludes that vessel. When that occurs, it is assumed no blood can enter the occluded vessels or any vessels downstream of the occluded vessel. The algorithm recalculates a baseline blood flow and resamples a clot to loop through the steps again.

If the clot reaches a terminal vessel, and it is still smaller than the vessel, it is assumed that the clot generates no occlusions in the penetrating vessels and enters the capillary bed. As we are trying to quantify the damage done by micro-emboli to the penetrating vessels, we do not create capillary bed occlusions in this simulation. The effect of micro-emboli on the capillary bed is thus considered separately in Section 4.9. As such, no vessels in the penetrating tree are occluded, and a new clot is sampled to loop through the algorithm again.

The algorithm described, shown in Fig 2, is repeated until only one penetrating vessel remains or until 50,000 clots have been sampled, whichever comes first. The large number of maximum clots is chosen such that we can ensure that all the vessels are eventually occluded, allowing us to track the damage done by micro-emboli sequentially.

### 4.7 Parameterising the penetrating arterioles

There are 2 specific parameters of interest when attempting to model the penetrating vessels over a large scale – the permeability and the coupling coefficients. The permeability of the penetrating vessels represents the ability of the trees to drive blood flow deeper into the grey cortex due to their 1-dimensional nature [54], and the coupling coefficients represent the pre-capillary arterioles’ ability to deliver blood to the capillary bed. These parameters are necessary to simulate large regions of the human brain blood flow efficiently [31]. However, these parameters also effectively average out the micro-scale fluctuations in the vasculature, replacing a complex network with one parameter. This can be problematic with the penetrating vessels when considering, for example, the terminal nodes of the trees. It is highly unlikely they are distributed evenly through the grey matter with depth. It is also highly unlikely that the permeability of the trees is constant with depth through the voxel. We mitigate this problem by splitting the voxel into 6 layers, similar to other approaches [37,55]. Permeabilities and coupling coefficients are then calculated for each of the 6 layers through the voxel, increasing the resolution of the large-scale simulations.

#### Permeability

As the permeability of interest is defined as the ability of the penetrating vessels to drive blood flow deeper into the cortex, it is calculated in the vertical direction only. The angle that every vessel makes with the vertical is calculated (situating the vertical at the vessel node where flow enters the vessel). The vertically projected flow of the vessel, *Q*_*Z,i*_, is then calculated as

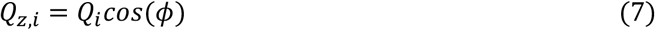

where *ϕ* is the angle between the vessel and the vertical, and *Q*_*i*_ is the absolute flow through the vessel. The arterial blood flow, *Q*_*Z,i*_, through each of the six layers is then summed separately and the permeability calculated as

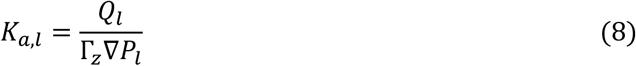

where *Q*_*l*_ = ∑ *Q*_*Z,i*_ through layer *l*, *K*_*a,l*_ is the arteriolar permeability in layer *l*, Γ_*Z*_ is the cross-sectional area through which the blood flows (in this case the area of the voxel), and ∇*P*_*l*_ is the pressure gradient across the layer. Using this method provides 6 different estimates of arteriolar permeability at 6 different depths in the grey matter.

#### Coupling coefficients

The method to calculate the coupling coefficients can be found in Hyde et al. [56]. The coupling coefficients can be thought of as a conductance (or 1/resistance) to flow from the arteriolar compartment to the capillary compartment via the pre-capillary arterioles. This can be written as

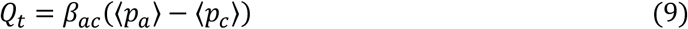

where *β*_*ac*_ is the coupling coefficient between the arteriolar and capillary compartments, 〈*p*_*i*_〉 is the volume averaged pressure of compartment *i*, and *Q*_*t*_ is the flow of the arterioles which terminate in the capillary compartment. This can be rewritten as

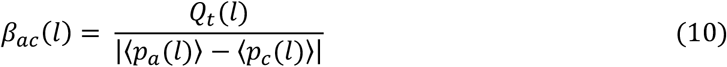

where the bracket *l* term refers to the values in layer *l*. The volume averaged flow in layer *l*, *Q*_*t*_(*l*), can be calculated using equation (1). The volume averaged pressure of the arteriolar compartment 〈*p*_*a*_(*l*)〉 is calculated via

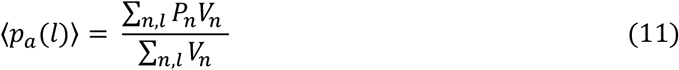

where *P*_*n*_ is the average nodal pressure of vessel *n* in layer *l*, and *V*_*n*_ is the volume of vessel *n* in layer *l*. Six values of average pressure are calculated, one for each layer.

Finally, the volume averaged capillary pressure at each layer 〈*p*_*c*_(*l*)〉 is calculated by taking the volume average of the pressures in the finite element mesh of the capillary bed. This allows for the permeability and coupling coefficients of the penetrating vessels to be fully characterised. In order to be able to characterise the variable permeabilities and coupling coefficients calculated above, a polynomial least squares fit is used to derive a function of how these coefficients vary with depth which is then fitted over the 100 simulations conducted on the statistically accurate voxels.

### 4.8 Investigating thrombectomy technique, clot composition, and individual microvasculatures

Based on the pipeline previously presented, we now seek to investigate whether the different micro-emboli distributions from thrombectomy have an impact on the permeability (and hence perfusion) of the penetrating arterioles following the introduction of micro-emboli. The micro-emboli algorithm is therefore used to simulate an emboli shower for 8 different scenarios – 4 techniques used on ‘hard’ or ‘soft’ clots.

As well as this, we investigate the impact that differing microvasculatures have on the resulting drops in permeability and coupling coefficients. Each of the 100 statistically accurate voxels can be thought of as an ‘individual’ microvasculature. We randomly choose 3 of these voxels and, for each voxel, we simulate 100 different sampled clot distributions entering the penetrating vessels. These simulations are compared to another simulation where we take our 100 voxels and run the same sampled clot distribution through each voxel – this is done for 3 different sampled clot distributions. The question that we seek to answer is whether it is the microvasculature or the clot distributions that primarily determine microvascular robustness to drops in perfusion. Differences between distributions are quantified using the correlation matrix distance (CMD) on the covariance matrices of each distribution [57].

### 4.9 Micro-occlusions in the capillary bed

Finally, the effect of micro-emboli on capillary beds is considered. Previously, we demonstrated how statistically accurate models of capillary beds can be homogenized into a porous medium [32]. This method is combined here with a vessel occlusion algorithm to simulate the effect of micro-emboli entering the capillary bed. The decrease in permeability as a function of the vessel fraction lost, surface area fraction lost, and volume fraction lost are all quantified. This allows for the scaling up of partially occluded capillary networks to large regions of tissue.

Details of the blood flow simulation and homogenization of the capillary bed can be found in El-Bouri & Payne [32]. In brief, Poiseuille flow is assumed through the capillary vessels, with the capillary network being periodic. A pressure gradient is imposed in one direction and the permeability – the ratio of the volume-averaged flow and the pressure gradient – is calculated. This is repeated for the 3 principal directions of the capillary network. The result is a 3×3 tensor, with 3 principal permeabilities (the permeability in the direction of the pressure gradient) and 6 cross-permeabilities.

As the smallest measurable clot was 8.034 μm, and the average diameter of a capillary is 6.23 ± 1.3 μm [40], the micro-emboli algorithm above could not be used on the networks as they would always occlude the inlet vessel. As such, a random vessel was chosen in the network and this vessel was fully occluded, with the permeability being recalculated after every occlusion. Three cube sizes were used to simulate the micro-occlusions with lengths of 375, 500, and 625 μm. Normalised permeability changes are calculated, and the effect of the micro-emboli on the capillary bed is quantified such that it can be used in full-brain simulations. Constrained least squares optimization is used to fit a line of best fit with the equation

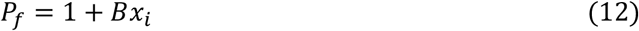

where *P*_*f*_ is the permeability fraction, *x*_*i*_ is the change in either vessel fraction, volume fraction, or surface area fraction of the networks, and *B* is the gradient. All parameters calculated for the capillary bed are based on 500 statistically accurate capillary networks at each cube size (1500 total).

## 5 Results

### 5.1 Healthy blood flow in the penetrating arterioles

Blood flow was simulated in 100 statistically accurate voxels of the cerebral microvasculature. The permeability and coupling coefficients were quantified for the penetrating arterioles in the ‘no-occlusion’ scenario. The permeability results are shown in Fig 3.

**Fig 3.**
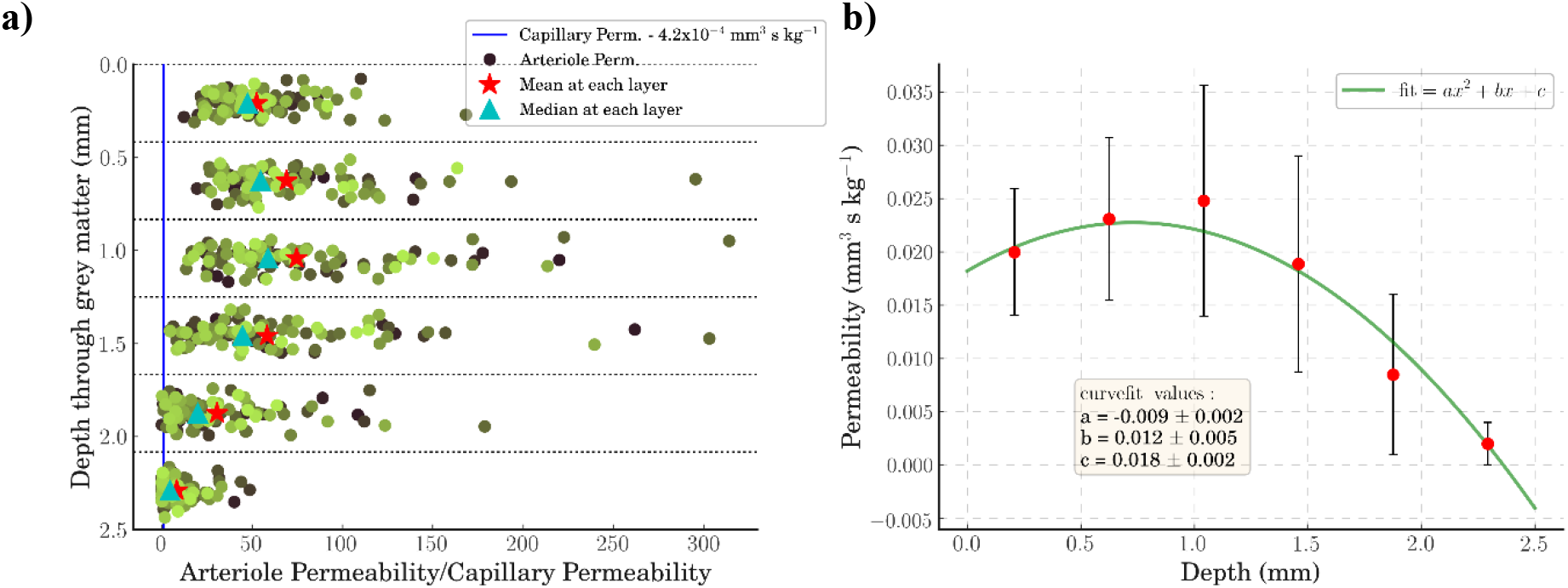
a) A scatter plot of the permeability of the 100 voxels at each of the 6 depth layers. The cyan triangle indicates the median permeability of the capillary bed. The mean arteriolar permeability is indicated with a red star at each layer. b) A quadratic line of best fit over the median permeabilities at each layer – error bars are interquartile ranges

The permeability increases with depth until the middle layers before dropping off towards the bottom layers, as not all penetrating vessels penetrate the full depth of the grey matter (Fig 3a). The mean value of the permeability across the layers is approximately 50 times the capillary permeability. At each layer the average permeability ranges from around 30x at the top layer, to 75x in layer 3, to 9x at the bottom layer. A quadratic curve fit is used to derive a function that models arteriolar permeability with depth, such that it can be used in future porous model implementations (Fig 3b). The permeability distribution at each layer can also be characterised as log-normal (Shapiro-Wilks test, *α* = 0.05) up to and including layer 4 (S1 Table).

The coupling coefficients demonstrate a similar drop from the top layers to the bottom layers, driven by the fact that there are fewer terminal nodes at the bottom layers [33]. The variability in the coupling coefficients from voxel-to-voxel is larger than the permeability variability (S1 Fig). The median values of the coupling coefficients are 2.8×10^−5^, 4.4×10^−5^, 4.3×10^−5^, 2.7×10^−5^, 1.1×10^−5^, and 4.3×10^−6^ Pa^−1^ s^−1^ (from layers 1 – 6). A quadratic curve-fit is used to derive a function that models the coupling coefficients with depth for future porous model implementations (S1 Fig). The coupling coefficient distribution can be characterised as log-normal (Shapiro-Wilks test, *α* = 0.05) up to and including layer 4 (S2 Table).

The above results parameterise the healthy penetrating vessels so they can be scaled up to large regions of the brain. More importantly, however, these healthy results provide a baseline from which the effect of micro-occlusions can be quantified.

### 5.2 The arterioles are robust to micro-emboli at the population level

As there are 4 thrombectomy techniques used with 2 clot consistencies to give 8 different micro-emboli distributions to choose from, a baseline must be chosen to conduct the following simulations. Here we choose the ADAPT technique with a hard clot to present the following results. A comparison with other techniques and clot consistencies can be found in Sections 5.4, 5.5.

The variable of interest here is the fractional drop in permeability. This is defined as the permeability divided by the initial healthy permeability. We compare this independent variable against 3 dependent variables – the fraction of vessels occluded, the fraction of vessel surface area occluded, and the fraction of vessel volume occluded. The surface area and volume drops have been calculated as these geometric parameters control oxygen transport in the microvasculature and can in future be used for validation. Fig 4 shows these results for 2 different layers – the top-most grey matter layer, and the middle layer (from 1.25 – 1.67 mm depth). An example of the permeability decrease against volume fraction decrease in all 6 layers can be found in S2 Fig.

**Fig 4.**
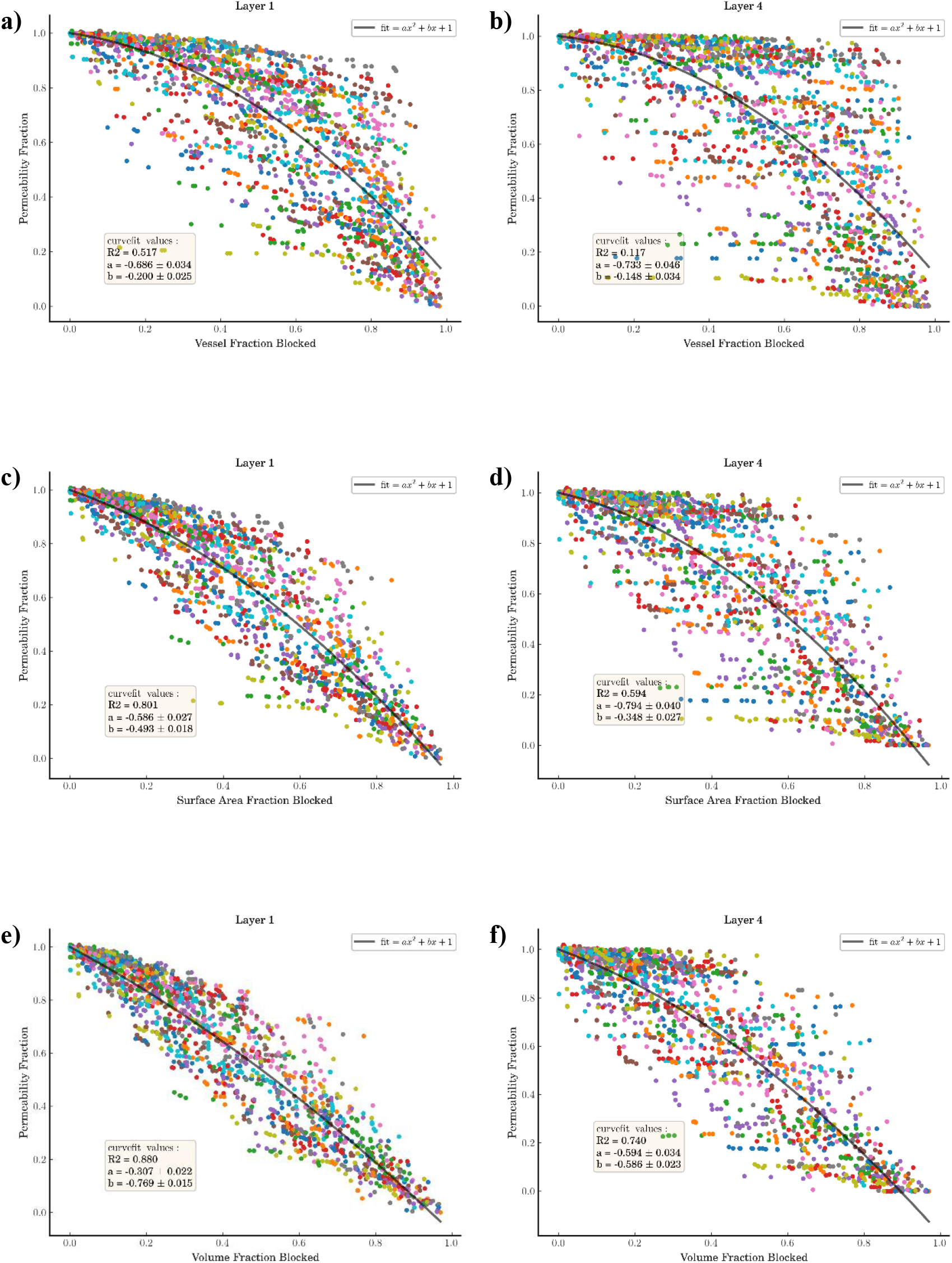
a,b) The fractional drop in permeability against the fraction of vessels blocked where a) is in the top layer of the voxel and b) is the middle layer. c,d) The fractional drop in permeability against the fraction of vessel surface area blocked where c) is in the top layer of the voxel and d) is the middle layer. e,f) The fractional drop in permeability against the fraction of vessel volume blocked where e) is in the top layer of the voxel and f) is the middle layer. The middle layer is at a depth of 1.25 – 1.67mm. A line of best fit is plotted in black, with the fit equation in the top right corner of each graph, with the bottom left of the graph giving the coefficients of the line of best fit and the R^2^ goodness-of-fit metric.

The permeability is generally robust with vessel fraction blocked, up to around 50 % of the vessels before the permeability starts to drop more rapidly. This can be explained by the fact that many small vessels are occluded first – due to 98% of all emboli having a diameter of < 20 μm for this configuration of clot consistency and thrombectomy technique – which has a relatively small impact on the permeability. On the other hand, the permeability drops almost linearly with volume fraction blocked with a 1:1 ratio in the top layer.

Interestingly, in the middle layer for all 3 dependent variables, the permeability’s robustness to occlusions appears to improve over the first 40-50% of vessel/volume occlusions. As the vessels continue to be occluded, however, the drop off in permeability is quicker than in the topmost layers. The variability increases with depth until by the time the bottom two layers are reached, there is almost a dichotomous choice between a permeability of 1 and 0 (S2 Fig). This is due to the sparse number of vessels in the lower layers, leading to them being vulnerable to very few micro-emboli.

The permeability, and its drops with successive micro-emboli, can thus be characterised as a 2-dimensional surface. The equation for the healthy permeability distribution with depth can be multiplied, at each depth layer, by the respective permeability drop equation for that depth layer. An example of a 2-dimensional surface generated with volume fraction as the dependent variable, ADAPT as the thrombectomy technique, and a hard clot can be found in Fig 5 (calculated using the healthy permeability equation from Fig 3b). These 2-dimensional functions can be calculated for all the different possible combinations of thrombectomy, clot, and dependent variable. This provides a simple look-up table to determine the change in permeability for a given technique, clot, depth, and fraction occluded.

**Fig 5.**
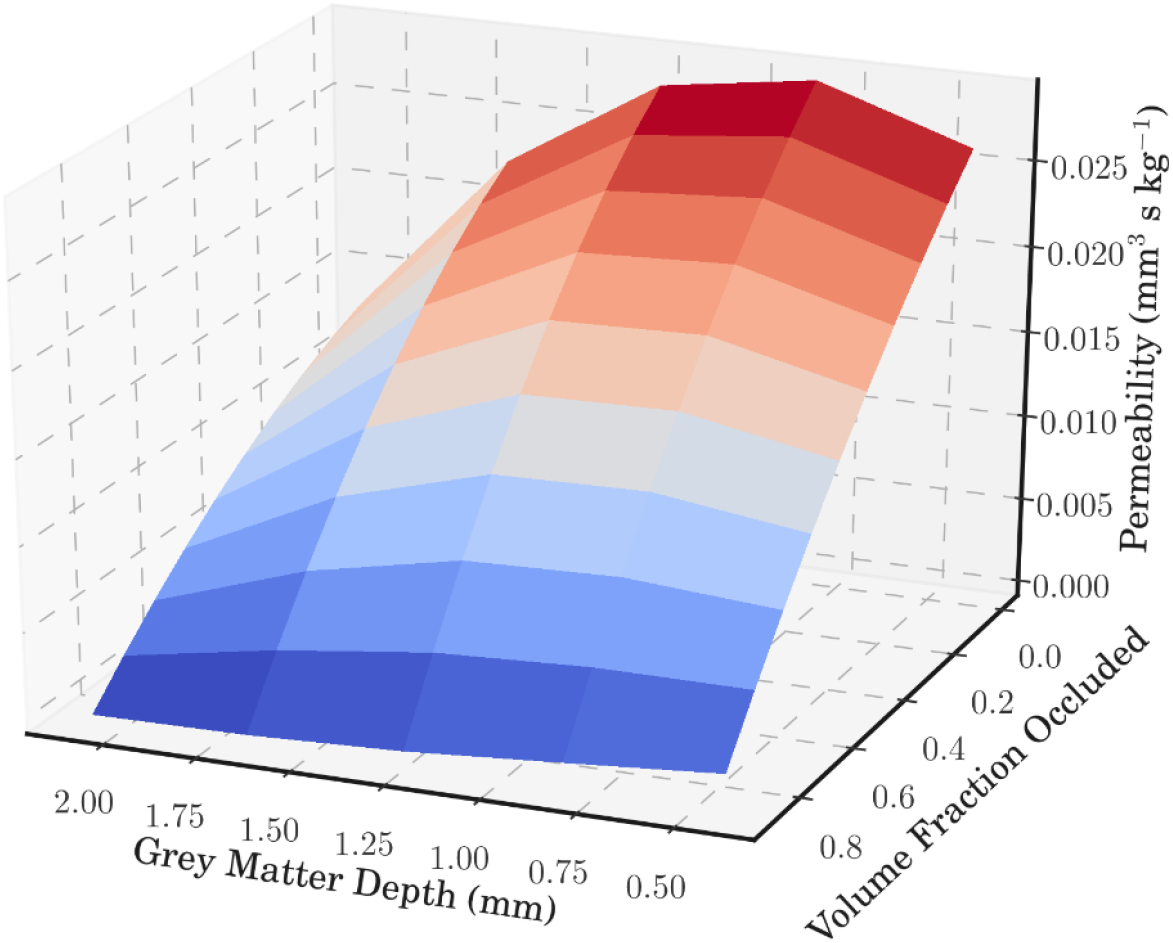
A 2-dimensional surface plot for the ADAPT thrombectomy technique removing a hard clot. The permeability is plotted against voxel depth and volume fraction of the vessels occluded.

The coupling coefficients also display non-linear behaviour with micro-emboli occlusions, but this behaviour is far more variable than the permeability results. This is due to blood being rerouted as occlusions occur, leading to coupling coefficients increasing as well as decreasing from the baseline. As well as this, the coupling coefficients are also dependent on the pressure in the capillary beds and terminal arterioles (Eq (10)). An example of the change in coupling coefficients with depth and volume occlusions can be found in S3 Fig.

### 5.3 The microvascular geometry dictates robustness to micro-emboli smaller than the penetrating vessel inlet diameter

To investigate the role of the microvasculature on robustness to micro-emboli, 3 voxels were chosen from our 100 voxels. Each voxel had 100 micro-emboli simulations with each simulation having a different clot distribution sampled. Fig 6 shows the changes of permeability for these 3 voxels for the middle-layer. As can be seen, it appears that the microvascular topology dictates the response to micro-emboli, with clear banded regions with non-linear jumps for some voxels, whilst other voxels have an almost linear drop with increasing occlusions.

**Fig 6.**
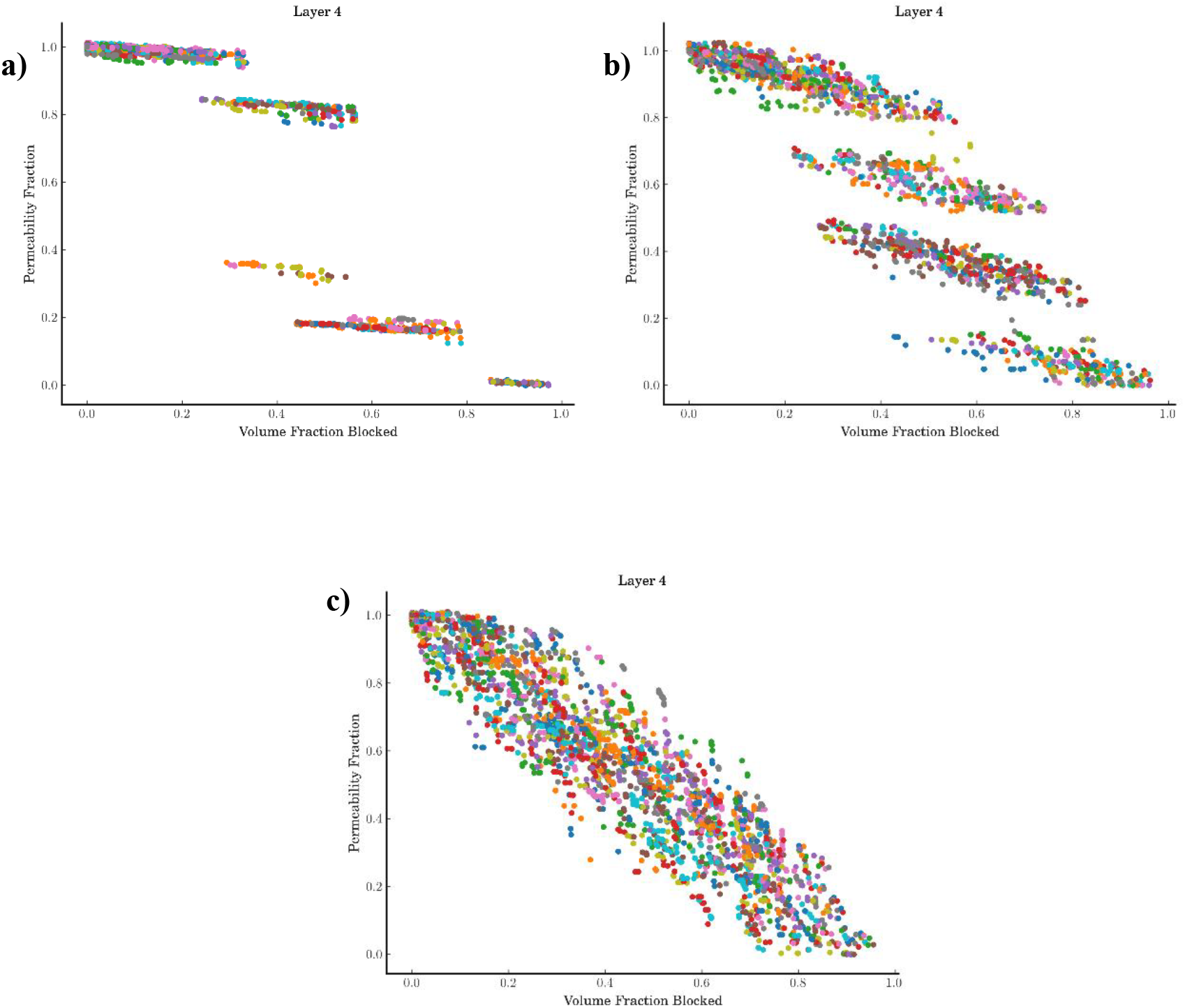
A comparison of the permeability drop against volume fraction occluded for 3 different voxel geometries simulated with 100 different clot distributions (for the ADAPT technique, hard clot) a) Voxel 1, b) Voxel 4, c) Voxel 31

In comparison, we simulated 100 different voxels but with the same sampled clot distribution entering each voxel (we fix the random seed sampling the clot distribution to be the same for each voxel). The aim of this was to investigate the effect of the sampled clot distribution variability on perfusion characteristics (similar to how we investigated the effect of microvascular variability in Fig 6). There was little visual difference found for the 3 sampled clot distributions (S4 Fig). Similar behaviour is also seen for the coupling coefficients, with linear drops in coupling coefficients seen for some voxels (S5 Fig).

Quantifying the statistical differences between these distributions was complicated by the bivariate nature of the distributions. As such, to provide a simple comparison, we use the correlation matrix difference (CMD) to compare the covariances of each distribution against the baseline distribution (Fig 4f). The CMD has a value of 0 when the covariance matrices are identical and a value of 1 when they are orthogonal. When comparing the 3 constant microvascular simulations the range of the CMD was 1×10^−4^ to 2.85×10^−3^. For the constant clot simulations, the range was 7×10^−6^ to 3×10^−4^ indicating that variability and robustness to micro-emboli is dominated by the microvascular geometry.

### 5.4 Hard clot micro-emboli lead to larger drops in permeability

The above analysis has all been conducted using ‘hard’ clot fragmentation data. We now simulate ‘soft’ clot fragmentation and the effect of these micro-emboli on flow characteristics. We are interested to observe whether clot consistency impacts downstream flow post-thrombectomy.

Visually, there is no discernible difference for the relationship between permeability and the dependent variable on the population level, but certain voxels display changes in their permeability relationship (S6 Fig). Comparing individual microvascular geometries in both hard and soft clot simulations, the CMD score ranges between 3×10^−4^ and 7×10^−3^, indicating a similar variability to that found between voxel comparisons. The results are similar for the coupling coefficients.

Regardless of whether there was a visual difference or not, all voxels with soft clot simulations required on average twice as many occluding clots to develop the same drops in perfusion as for the hard clots. As well as this, around 4.5% of the clots sampled from the hard clot distributions went into occluding penetrating vessels (the rest passed through the arterioles into the capillary bed). For the soft clots, this was only 2.5% (Table 1). Clearly, for a given number of micro-emboli the smaller clot fragments of the soft clot lead to smaller drops in permeability.

**Table 1.** Summary of the differences between thrombectomy technique and clot consistency when considering total micro-emboli sampled and those micro-emboli that occlude vessels. The values are averaged over the 100 voxel simulations. The percentage of sampled clots occluding vessels is the average of the 100 individual percentages.

| *Thrombectomy Technique* | *Clot Consistency* | Average number<br>micro-emboli<br>sampled | Average number<br>micro-emboli<br>occluding vessels | Percentage of<br>sampled clots<br>occluding vessels<br>(%) |
| --- | --- | --- | --- | --- |
| *ADAPT* | *Hard* | 616.1 | 24.3 | 4.5 |
| *ADAPT* | *Soft* | 2704 | 48.1 | 2.5 |
| *BGC* | *Hard* | 208.5 | 25.8 | 14.0 |
| *BGC* | *Soft* | 994 | 26.4 | 3.0 |
| *GC at cervical ICA* | *Hard* | 1152.7 | 29.0 | 2.9 |
| *GC at cervical ICA* | *Soft* | 1146.6 | 22.9 | 2.3 |
| *Solumbra* | *Hard* | 76.4 | 18.6 | 26.3 |
| *Solumbra* | *Soft* | 1720.2 | 36.3 | 2.6 |

### 5.5 Thrombectomy technique has a large impact on clot fragmentation and downstream flow characteristics

The last variable of interest here is the thrombectomy technique. In the in-vitro experiments, 4 techniques were used; these are abbreviated as ADAPT, BGC, GC at cervical ICA, and Solumbra [20]. The emboli distributions from these 4 techniques used on hard and soft clots were used to simulate occlusions in the penetrating vessels. The results are shown in Fig 7 where the scatter plots have been binned into deciles with mean and standard deviation for ease of viewing.

**Fig 7.**
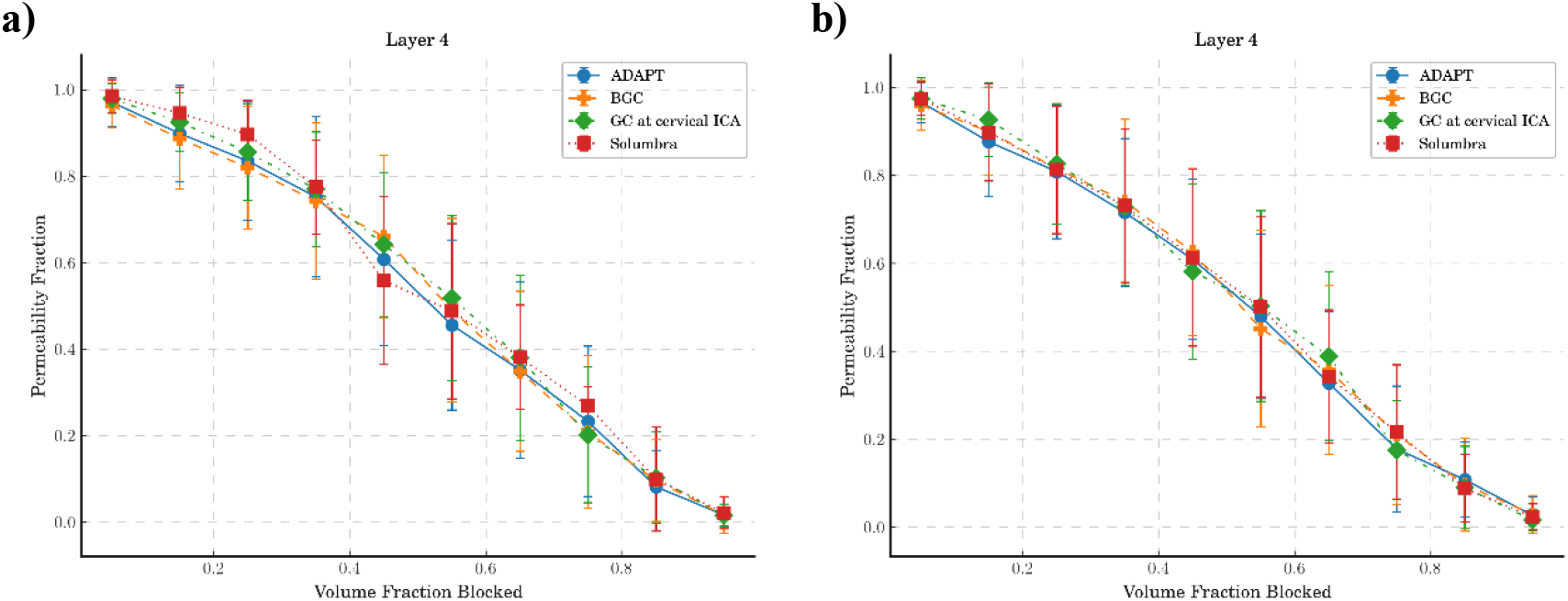
Comparison of the permeability drops against volume fraction blocked in the middle layer for 4 different thrombectomy techniques: a) removing hard clots and b) removing soft clots

Despite there being little visual difference between the 4 techniques, when analysing the numbers of clots required to reach similar drops in permeability, large differences emerge (summarised in Table 1). We measured: the average number of micro-emboli sampled in total to fully occlude the voxel (column 3 of Table 1); the average number of these micro-emboli that occlude a penetrating vessel (column 4 of Table 1); and hence the percentage of sampled clots that occluded the penetrating vessels (column 5 of Table 1). Simulating the Solumbra technique on a hard clot resulted in the fewest clots required to fully occlude the voxel; on average over the 100 voxels 18.6 clots (and hence 26% of sampled clots) sent through the penetrating vessels were required to fully occlude the voxel. The BGC technique had the next fewest clots required with 26 clots and 14% of the sampled clots. The GC at cervical ICA technique, on the other hand required 29 clots and 2.9% of the sampled clots.

In contrast, when using the soft clots, the numbers of clots required to achieve similar permeability drops drastically increased – except for GC at cervical ICA which had similar results compared to when using the hard clots. Both the Solumbra and ADAPT techniques saw a doubling of the clots required.

Clearly, there is a trade-off between the number of emboli that fragment off a clot, the size of these emboli, and the impact they have on downstream flow. The BGC and Solumbra techniques were previously reported to minimise the fragmentation of hard clots [20], but this also means that these clots are larger and hence more prone to occlude larger vessels (as indicated by our large percentage of clots that occlude vessels in the above results). On the other hand, the ADAPT and Solumbra techniques used on soft clots lead to twice as many clots being required to occlude the penetrating vessels to achieve similar drops in permeability.

### 5.6 Occlusions in the capillary bed lead to large drops in permeability

Finally, we simulated micro-emboli occluding capillary bed networks. These were simulated for 500 statistically accurate networks at 3 different cube length sizes (375 μm, 500 μm, and 625 μm). As the results were not statistically different over the 3 sizes, only the 375 μm results are presented in Fig 8. The permeability maintained its isotropy in the presence of micro-emboli hence it is presented here as a scalar.

**Fig 8.**
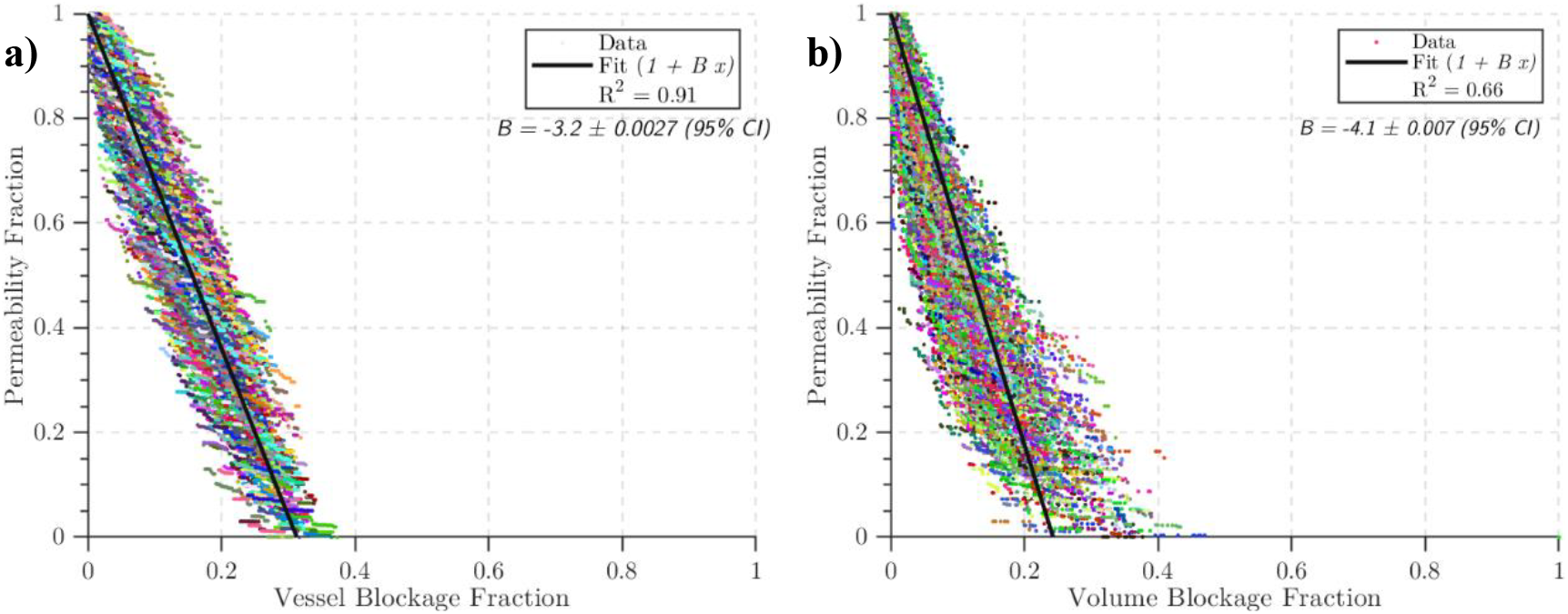
The drops in permeability over 500 statistically accurate capillary networks with a cube length of 375 μm. Lines of best fit are in black, with the gradient *B* in the top right corner of each graph. a) The fractional drop in permeability with fraction of vessels occluded (*B* = −3. 2), b) the fractional drop in permeability with vessel volume occluded (*B* = −4.1).

The permeability drop in the capillary bed was large in comparison to the drop in the penetrating vessels: −8 % / % surface area lost, −3.2 % / % vessel blocked, and −4.1 % / % volume blocked (see Fig 8). This is in line with a previous study where a −2.5 % drop in CBF/ % vessels blocked was found in simulations of occlusions in mouse and human vascular networks, as well as mouse synthetic networks [13]. It should be noted that the synthetic networks used in [13] had uniform diameter and length, unlike the networks in this paper which have length and diameter distributions that match physiological distributions.

Assuming a micro-emboli shower is spread evenly over the MCA territory (approx. 100 mL of tissue), 1.5% of this volume is capillary vessels, and a vessel density of 8000 vessels/mm^3^, around 120,000 micro-emboli would have to enter the capillary bed to cause a 4% drop in permeability (note that this is an overestimate as micro-emboli are not likely to spread evenly over the brain tissue). Depending on thrombectomy technique, the number of micro-emboli fragmenting off the clot can vary from a few thousand to over 500,000 [20]. It therefore appears that if micro-emboli enter the capillary bed there is significant potential for damage and large drops in perfusion.

## 6 Discussion

We return to the questions posed at the end of the introduction. First, can the changes in the blood flow modelling parameters due to micro-emboli showers be accurately quantified such that they can be used in the full organ models? Using a combination of in-vitro experiments of clot fragmentation and in-silico blood flow modelling, we have developed simulations of micro-emboli occluding vessels in the penetrating arterioles and capillary bed. Using these models we characterised ‘averaged’ flow parameters, such as permeability and coupling coefficients, in healthy conditions and under micro-emboli showers, for a range of thrombectomy techniques and clot consistencies. This now enables us to simulate the impact of these micro-emboli showers – after a thrombectomy – on the downstream tissue and its potential contribution to the ‘no-reperfusion’ phenomenon.

Second, what impact do thrombectomy technique and clot consistency have on downstream blood flow post-thrombectomy? Clot consistency and thrombectomy techniques have been shown here to have a large impact on the blood flow parameters. Whilst the relationship between permeability and volume of vessel occluded was similar, the numbers of micro-emboli required were substantially different, with soft clots on average requiring twice as many micro-emboli to reach similar drops in permeability. The Solumbra and BGC techniques used on hard clots resulted in larger micro-emboli entering the microvasculature, and hence fewer vessels needing to be occluded to reach large drops in permeability. Therefore, whereas the Solumbra and BGC techniques result in fewer emboli fragmenting off the clot [20], these emboli can cause more damage to the microvasculature due to their larger sizes.

In addition to this we investigated the impact of microvascular morphometry on blood flow parameters. Microvascular morphometry determined the robustness of penetrating vessels to occlusions, not the sampled micro-clot distribution. Therefore, if we can measure the microvasculature, even if only indirectly, we can determine whether the microvasculature will be robust to a micro-emboli shower. Characterising the microvasculature and its robustness, e.g. through the residue function [34], will be the subject of future work.

Finally, we compared drops in permeability for the penetrating vessels and capillary bed. Whilst the penetrating vessels had an approximately 1:1 relationship between volume lost and permeability drop, the capillary bed lost 4% of permeability per 1% volume drop. This lack of robustness to occlusions in the capillary bed is surprising but has previously been reported elsewhere with capillary stalling in mouse capillary beds. In blood flow simulations, Cruz-Hernandez et al. found that for every 1% of vessels stalled, the flow drops by 2.5% [13]. Our results indicate a 3.2% drop in permeability (a surrogate for flow) for every 1% of vessels blocked, which show good agreement for early studies. Permeability is a more robust metric than CBF here due to CBF being dependent on the volume of tissue simulated. As well as this, Schmid et al. found that the severity of a micro-occlusion in the capillary bed depends on the connectivity and original flow rate of the occluded vessel, with large reductions in flow for specific types of connectivities [29]. As we found for the penetrating vessels, morphometry of the microvasculature plays a large role in robustness to micro-occlusions.

The smallest micro-emboli sizes available from our experimental data were 8 μm. While this was sufficient for micro-emboli occlusions in the penetrating vessels, we did not have information on smaller micro-emboli that would occlude the capillary vessels. This meant we could not track emboli through the capillary bed. In future, data on micro-emboli < 8 μm in diameter would help in more clearly understanding how micro-emboli affect the capillary bed. The diameters measured were the Feret diameters of the micro-emboli, however we used these diameters to assume spherical micro-emboli. It is therefore possible that our drops in permeability are over-estimated, as the clot could orient along a smaller dimension and pass through the vessel.

It should be noted that a limitation of the analysis presented here is the separate treatment of the penetrating arterioles and capillary bed. This was done for two reasons. Firstly, to provide a comparison to previous studies, as discussed above. Primarily, however, this was done to fit into our full brain modelling framework where the capillary bed and penetrating arterioles are treated as separate compartments coupled together via coupling coefficients. The effects of extravasation of the micro-emboli in the capillary bed have also been neglected, and these will likely result in restoration of flow over several days [28]. This model is also purely passive, with no hyperaemia or regulation of vascular tone, which likely impact passage of micro-emboli through the microvasculature.

The final question posed was: To what extent is clot fragmentation responsible for the post-thrombectomy no-reflow phenomenon? We only analysed the impact of micro-emboli in voxels, or cortical columns of microvasculature. In order to fully understand the impact of micro-emboli on the vasculature, a full brain simulation is required. In future, we will use the work developed here to simulate micro-emboli showers in a full-brain in-silico simulation of thrombectomy. Blood flow parameters in the full-brain model will be updated using the parameters derived here, and the impact of micro-emboli in a region of the brain can be compared and validated against CT perfusion and MR scans of patients pre- and post-thrombectomy. Furthermore, we will investigate further the impact of microvascular morphometry on blood flow drops and how this can potentially be used to infer patient robustness to micro-emboli.

## 8 Supporting Information

**S1 Table. The main statistics for the distribution of permeabilities in each layer of the healthy voxels (for 100 voxels).** The first 4 layers can be approximated as log-normal. The p-value is the Mann-Whitney U-test between adjacent layers to determine if adjacent layers have similar distributions of permeability. For comparison, the capillary bed permeability is 4.28×10^−4^ mm^3^ s kg^−1^

**S2 Table. The main statistics for the distribution of coupling coefficients in each layer of the healthy voxels (for 100 voxels).** The first 4 layers can be approximated as log-normal. The p-value is the Mann-Whitney U-test between adjacent layers to determine if adjacent layers have similar distributions of coupling coefficients.

**S1 Fig. a) A scatter plot of the coupling coefficients of the 100 voxels at each of the 6 depth layers. The mean arteriolar coupling coefficient is indicated with a red star at each layer. b) A quadratic line of best fit over the median coupling coefficients at each layer – error bars are interquartile ranges**

**S2 Fig. a-f) The drop in permeability with respect to volume fraction blocked over the 6 layers, starting with the top layer a) and ending at the bottom layer f).**

**S3 Fig. a-f) The change in coupling coefficients with respect to volume fraction blocked over the 6 layers, starting with the top layer a) and ending at the bottom layer f)**

**S4 Fig. Permeability drops for 3 different sampled clot distributions over 100 voxel geometries. Results are shown for the middle layer in the voxels.**

**S5 Fig. A comparison of the coupling coefficient drop against volume fraction occluded for 3 different voxel geometries simulated with 100 different clot distributions (for the ADAPT technique, hard clot) a) Voxel 1, b) Voxel 4, c) Voxel 31**

**S6 Fig. A comparison of the effect of hard and soft clots on permeability drops with respect to volume fraction occluded a) Hard clot for all 100 voxels b) Soft clot for all 100 voxels, c) Hard clot for Voxel 1 simulated with 100 clot distributions, d) Soft clot for Voxel 1 simulated with 100 clot distributions, e) Hard clot for Voxel 4 simulated with 100 clot distributions, f) Soft clot for voxel 4 simulated with 100 clot distributions**

